# Insights into Long-term Acclimation Strategies of Grapevines in Response to Multi-decadal Cyclical Drought

**DOI:** 10.1101/2022.05.05.490818

**Authors:** Dilrukshi S. K. Nagahatenna, Tarita S. Furlan, Everard J. Edwards, Sunita A. Ramesh, Vinay Pagay

## Abstract

The Australian wine industry is currently under pressure to sustain its profitability due to climate change. Therefore, there is a pressing need to explore grapevine genetic diversity and identify superior clones with improved drought resistance. We previously characterised more than 15,000 dry-farmed (for over 65 years) Cabernet Sauvignon clones in a vineyard and identified three drought-tolerant (DT) clones, which can maintain significantly higher intrinsic water use efficiency (*WUE*_*i*_) under limited soil moisture than drought-sensitive (DS) clones. To understand whether DT clones grown under multi-decadal cyclical drought can prime their vegetatively-propagated clonal progenies for future drought events, in this study, all DT and DS vegetative progenies were propagated with commercial clones in the glasshouse. Their physiological and molecular responses were investigated under well-watered and two recurrent drought (D1 and D2) conditions. We observed that concentration of a natural priming agent, γ-amino butyric acid (GABA), were significantly higher in all DT progenies relative to other progenies under drought. Both commercial and DT progenies exhibited improved gas exchange, photosynthetic performance and *WUE*_*i*_ under recurrent drought events relative to DS progenies. Our results suggest that DT progenies have adapted to be in a “primed state” to withstand future drought events.

## Introduction

The viticulture industry, which consists of wine, raisin and table grape production is the largest fruit industry in Australia. Currently, Australia is the world’s fifth-largest wine exporter and in 2019, the wine sector contributed $45.4 bn to the Australian economy [1]. Due to recent extreme drought events, heat waves and bushfires, Australian grapevine production reduced 20% in 2020, and the smallest vintage was recorded: 1.4–1.5 M tons compared to the average 1.75 M tons [2]. Given that the incidence and severity of drought events are predicted to increase in the future [3], there is an increasing need to select and/or breed superior grapevine cultivars and clones, which could perform better in dry climates.

Grapevine planting materials were first introduced to Australia from Europe in 1832. There was a massive expansion of genetic resources in Australia in 1960s due to mass introduction of Phylloxera-free clones with superior agronomic and oenological performances from overseas [4]. A clone is referred to as a vegetatively propagated population of vines originated from a single parental vine and clonal progenies are generally considered to be genetically identical to their parental vine. However, over time diverse notable phenotypic variations can emerge due to progressive accumulation of spontaneous mutations [5] or epigenetic modifications [6,7]. Since the beginning of the viticulture in Australia, clonal selection was employed as the main tool in grapevine improvement programs to preserve elite clones with desirable characteristics [8,9]. Australian old vineyards still preserve extensive genetic diversity of European selections, but intravarietal genetic variability has not been completely explored in Australia [9]. Therefore, exploring the superior genotypes hidden within early introductions would pave the wave for improving complex traits such as drought tolerance and water use efficiency (WUE).

Drought tolerance is a highly complex trait, which is determined by genotype and environmental interactions [10-12]. Although over the past decades, our knowledge on short-term drought acclimation responses has increased, a comprehensive picture of how key plant physiological processes are regulated under prolong cyclical drought episodes is largely unknown. Stomatal closure is one of the earlier responses to water deficit. It has long been known that root-derived stress signalling molecules such as abscisic acid (ABA) sense subtle changes of soil moisture and transduce the signal to upper parts of the plant for inducing stomatal closure [13-16]. However, there is still controversy regarding the proposed role of ABA [17]. Recent studies have demonstrated that aquaporin (AQP)-mediated hydraulic signals and chemical signals such as γ-amino butyric acid (GABA) and CLAVATA3/embryo-surrounding region-related (CLE) small peptides also contribute to long-distance communications in response to drought stress [18,19]. Stomatal closure is crucial in preventing excessive transpirational water loss, however it dramatically reduces photosynthesis due to limitation in CO_2_ influx [20]. When field-grown grapevines undergo severe water stress, photosynthesis is further constrained by reduction in mesophyll conductance (*g*_*m*_), biochemical limitations and reactive oxygen species (ROS)-mediated photooxidative damage [21,22]. A considerable body of evidence has recently indicated that prolonged exposure of a plant to mild to moderate stress conditions can effectively stimulate faster and stronger tolerance to subsequent stress events through the acquisition of a “stress memory” [23-25]. Interestingly, in some instances, priming-induced “stress memory” has shown to be inherited to seed-derived offspring [26,27]. Recent work has discovered certain priming elicitors that can trigger natural defence mechanisms in a metabolically cost-effective manner. For instance, several studies have shown that plants accumulate γ-amino butyric acid (GABA), a non-protein amino acid, at the onset of the stress [28,29]. GABA is a metabolite and stress signalling molecule which is synthesized from glutamate in the cytosol [30,31] and metabolised through the GABA shunt pathway in both cytosol and mitochondria [32]. Increased concentrations of GABA under drought stress has shown to induce stomatal closure via activation of anion channel, aluminium-activated malate transporters (ALMTs) [33]. Accumulation of GABA under stress conditions also helps activating plant’s innate defence potential and pre-conditioning of the plant to next drought event through synthesising osmolytes [34], enhancing photochemical efficiency and WUE and ROS detoxification [35-37].

The overarching aim of this study was to understand whether woody perennial crops such as grapevine grown under multi-decadal cyclical drought can prime their vegetatively propagated clonal progenies for future drought events. As the first step towards understanding this, we characterised more than 15,000 dry-farmed Cabernet Sauvignon clones that were planted in 1954 as a mass selection of unknown clones in a South Australian vineyard. Due to the extended period of dry-farming, over 65 years, these vines are ideal for unravelling long-term drought adaptation mechanisms. In our pilot field trial, we identified several drought-tolerant (DT) clones which can maintain significantly higher intrinsic water use efficiency (*WUE*_*i*_) under limited soil moisture and drought-sensitive (DS) clones [38]. We hypothesized that these superior DT clones may have already primed due to long-term exposure to limited soil moisture and thereby may have the ability to successfully transmit drought acclimation strategies to their subsequent clonal progenies. In this study, we examined this hypothesis by evaluating the drought acclimation responses of their clonal progenies under two recurrent cyclic drought events in the glasshouse. To the best of our knowledge, the results obtained here are the first demonstration which provides novel insights into transgenerational and intragenerational drought priming mechanisms in grapevine clones.

## Materials and Methods

### Plant Materials

The drought-tolerant (DT) and drought-sensitive (DS) clones were selected based on differences in *WUE*_*i*_ observed under field conditions as described in Pagay *et al*. (2022) [38]. Five cuttings each were vegetatively propagated from previously selected three drought-tolerant (DT) and two drought-sensitive (DS) dry-farmed clones along with three commercial clones (G9V3, CW44 and SA125) at the Yalumba Nursery (Nuriootpa, SA, Australia). Propagated vines were re-potted in 4.5 L pots with a mixture of 50% University of California soil Mix (61.5 L of sand, 38.5 L of peat moss, 50g of calcium hydroxide, 90g of calcium carbonate, and 100g of Nitrophoska (12:5:1, N: P: K plus trace elements) and 50% perlite and vermiculite mix (50:50) at the Plant Research Centre (Waite Campus, University of Adelaide, SA, Australia). In order to facilitate similar growth rates, all vines were pruned to four nodes and incubated in a dark cool room at 4°C for at least 25 days. All clones were grown under the glasshouse conditions with 16 h photoperiod. Two overhead supplemental light sources were turned on from 6:00 to 20:00 every day to maintain uniform light distribution and intensity independently of external environmental conditions. Temperature and relative humidity were continuously recorded using data loggers (Tinytag Plus 2, Gemini, UK) in the glasshouse. After budburst, the vines were treated with 1.6 mL/L of Megamix® (13:10:15 N: P: K plus trace elements) every second week. When they reached 20-leaf stage, they were pruned to a similar leaf area.

### Drought cycles

In order to compare drought stress responses between pre-selected DT and DS clones, all 15 vines from 3 DT clones were grouped and compared with DS clonal group which consisted of 10 vines and commercial clonal group with 15 vines. Each group will be mentioned as DT, DS and commercial clonal progenies hereafter. Nine vines from each DT and commercial clonal progenies and 6 from DT progeny (3 vines/clone) were subjected to two cyclic drought events (treatment group) and remaining vines (6 from DT and commercial and 4 from DS progenies) were included in the well-watered treatment (control group). All vines were randomly positioned within each group. Vines in the control group were watered daily to achieve an equivalent pot weight. Soil moisture was measured daily using a Teros-10 soil triple sensor (METER, WA, USA). For evaluating clonal performances upon drought stress, water supply was withheld for the treatment group until soil moisture reached 4% volumetric soil water content (VSW) or less. If the soil moisture reduced under 4%, the vines were watered to the equivalent pot weight until the drought cycle was finished. Then they were rewatered to field capacity and until stomatal conductance retuned to similar values displayed by control vines (Figure 1a). After full recovery from D1 (21 days after rewatering), second drought cycle was imposed by withholding water as described above. Measurements and leaf tissue sampling were taken before drought stress (Day 0), at the peak of the 1^st^ drought cycle (7 days after withholding water-D1), after full recovery from D1 (21 days after rewatering (day 0 of the second drought cycle), and at the peak of the 2^nd^ drought cycle (4 days after withholding water-D2).

**Figure 1.**
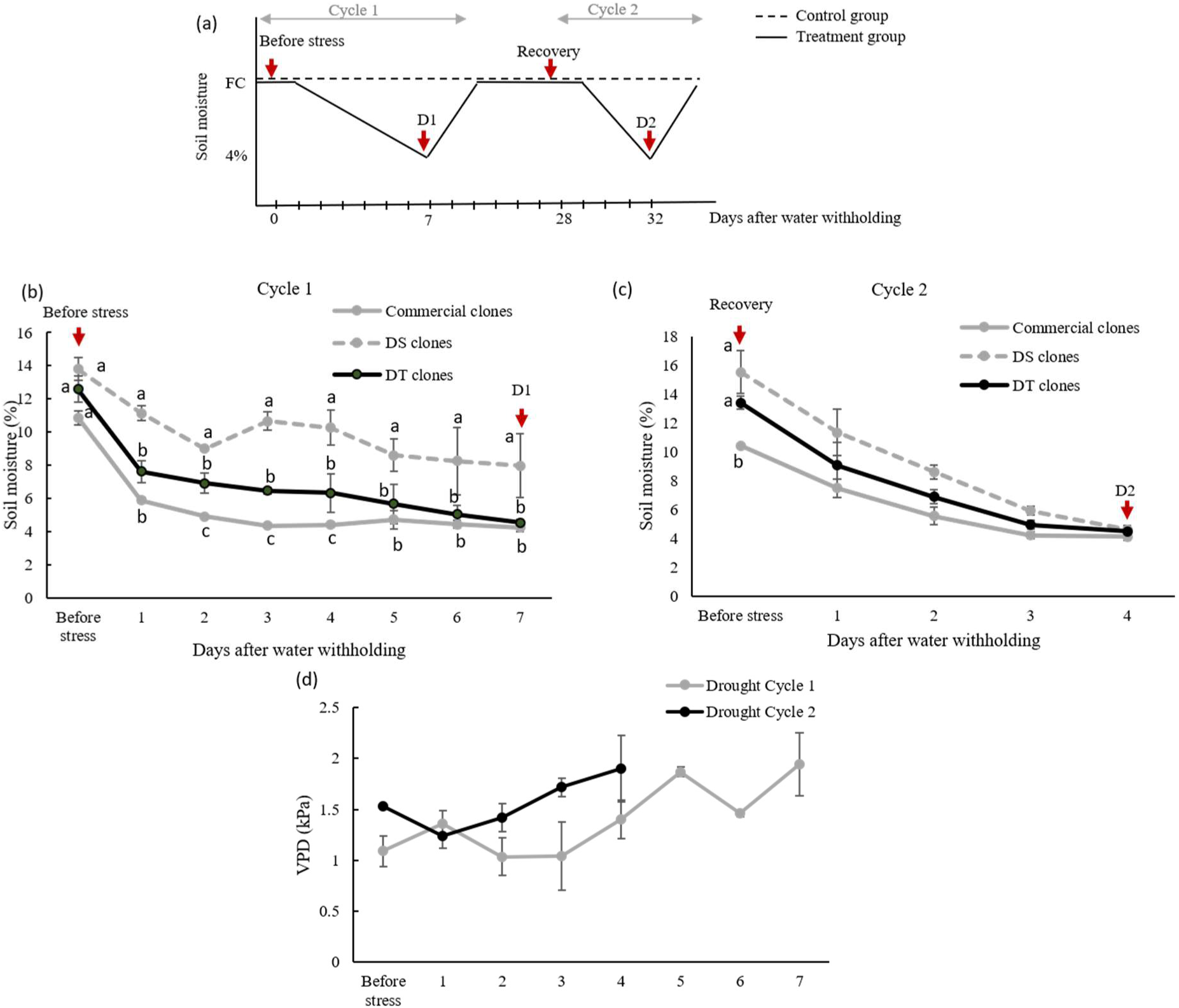
(a) Schematic representation of the experimental design: variation in soil moisture in both control (dotted lines) and treatment groups (solid lines) during two cycles of dehydration and rehydration. Arrows indicate four different time points where measurements were taken (before stress, the peak of the 1^st^ drought cycle (D1), after full recovery, and the peak of the 2^nd^ drought cycle (D2)). (b) Depletion of soil moisture in drought-treated commercial, DS and DT clones during 1^st^ and (c) 2^nd^ dehydration cycles, and (d) changes in vapour pressure deficit (VPD) during 1^st^ (grey line) and 2^nd^ (black line) drought events. Values are means ± SEM of six to nine biological replicates. Statistical analysis was conducted using two-way ANOVA and lowercase letters denote statistically significant differences (*P*<0.05) between clones at each time point.

### Midday stem water potential (Ψ_s_) and leaf Gas exchange measurements

Midday stem water potential and *in vivo* gas exchange parameters were measured using a Scholander-type pressure chamber (Model 1505, PMS Instruments, Albany, NY USA) and LI-6400XT (LI-COR Inc., Lincoln, NE, USA) respectively as described in Pagay *et al*. (2022) [38].

### Chlorophyll fluorescence

Chlorophyll fluorescence parameters were measured with a LI-6400XT equipped with a leaf chamber fluorometer (model: 6400-40, LI-COR Inc., Lincoln, NE, USA) in dark- and light-adapted fully expanded leaves positioned in the bottom, middle and top levels per vine. In light-adapted leaves, the steady-state fluorescence yield (_*s*_) was measured. Then a saturating white light pulse (8000 μmol m^-2^ s^-1^) was applied for 0.8 s to achieve the light-adapted maximum fluorescence (*F*_*m*_′). The actinic light was then turned off, and far-red illumination was applied (2 μmol m^−2^ s^−1^) to measure the light-adapted initial fluorescence (*F*_*0*_′). Vines were kept overnight in darkness for dark-adapted measurements and basal fluorescence (*F*_*0*_) and maximum fluorescence emission (*F*_*m*_) were measured by illuminating leaves to weak modulating beam at 0.03 μmol m^-2^s^-1^ and saturating white light pulses of 8000 μmol m^-2^ s^-1^, respectively. Non-photochemical quenching (*NPQ*), photochemical quenching coefficient (*qP*), and actual photochemical efficiency of PSII (Φ_PSII_) were calculated as: *NPQ* = (*F*_*m*_−*F*_*m*_*′*) / *F*_*m*_*′* [39], *qP* = (*F*_*m*_*′*– *F*_*s*_)/(*F*_*m*_*′*– *F*_*0*_*′*) [40], Φ_PSII_ =(*F*_*m*_*′*–*F*_*s*_)/*F*_*m*_*′* [39]. Electron transport rate (ETR) was calculated according to Rahimzadeh-Bajgiran, *et al*. [41] using the following equations.

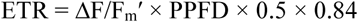

Where, ΔF/F_m_′ is the PSII photochemical efficiency, PPFD is the photosynthetic photon flux density incident on the leaf, 0.5 is a factor that assumes equal distribution of energy between the two photosystems, and 0.84 is the assumed leaf absorptance [42,43]. Mesophyll conductance (*g*_*m*_) was calculated using a theoretical value of non-respiratory compensation point G*=42.9 μmol mol^-1^ (ppm) CO_2_ [44].

### Abscisic Acid (ABA) quantification

Approximately 30 μl of Xylem sap was extracted from leaves at the end of *Ψ*_*s*_ measurements by increasing the balancing pressure by 0.2-0.4 MPa. Sap samples were snap frozen in liquid nitrogen and subsequently transferred to a −80°C freezer until analysis. ABA abundance in xylem sap (ABA_xyl_) was analysed by liquid chromatography/mass spectrometry (LC MS/MS, Agilent 6410) [12].

### Analysis of Ascorbate peroxidase (APX) enzyme activity

0.5 g of frozen leaf samples were suspended in 2 ml of 0.1 M sodium phosphate buffer (pH 7.0) and incubated for 10 min on ice. Samples were centrifuged at 12,000 g for 15 min and 10 µL of the supernatant from each sample was mixed with 290 µL of assay mixture consisting of 0.5 mM ascorbic acid, 0.1 mM EDTA-Na_2_ and 0.1 mM H_2_O_2_ solutions prepared in 0.05 M sodium phosphate buffer (pH 7.0). The APX activity was determined by measuring the absorbance at 290 nm, in a FLUOstar Omega plate reader (BMG LABTECH GmbH, Ortenbery, Germany). The decrease in absorbance corresponded to oxidation of ascorbic acid. One enzyme unit was defined as 1 mol of ascorbic acid oxidized per minute at 290 nm [45].

### Quantification of endogenous GABA concentrations

Leaf samples were snap frozen in liquid nitrogen and 0.1g of frozen ground samples were used for GABA extraction. GABA quantification was conducted according to the protocol described in Ramesh, *et al*. [46].

### Gene expression analysis by Quantitative real-time PCR

Grapevine leaves that were used in *Ψ*_*s*_ measurements were snap frozen and stored at −80 ^0^C freezer. RNA was extracted using Spectrum plant total RNA kit (Sigma, USA) according to the manufacturer’s instructions, and contaminated DNA was removed according to On-column DNase digestion protocol (Sigma, USA). Total RNA was quantified with a UV spectrophotometer and quality of RNA was assessed by gel-electrophoresis. For cDNA synthesis, 1 μg of total RNA was reverse transcribed using the iScript cDNA synthesis kit (Bio-Rad, CA, USA). In Quantitative real-time PCR (qPCR), 1 μL of 1/10 diluted cDNA was amplified in a reaction containing, 5 μL KAPA SYBR® FAST Master Mix (2X) Universal (Kapa Biosystems Inc., MA, USA), and 100 nM of gene-specific primers. qPCR primer sequences for two aquaporin (AQP) genes (*VvTIP2;1* and *VvPIP1;1*) and two stable housekeeping genes, *VvELF* and *VvUbi* were obtained from Shelden, *et al*. [47]. The amplification was conducted in a QuantStudio 12K Flex Real-Time PCR system (ThermoFisher Scientific, USA) according to the following conditions: one cycle of 3 min at 95 °C followed by 40 cycles of 16 s at 95 °C, and 20 s at 60 °C. To ensure single-product amplification, melt curve analysis was performed by heating the PCR products from 60°C to 95 °C at a ramp rate of 0.05 °C s^-1^. A two-round normalization of qPCR data was carried out by geometric averaging of multiple control genes as described by Vandesompele, *et al*. [48] and Burton, *et al*. [49].

### Statistical analysis

In order to understand the long-term drought acclimation responses of dry-farmed clones, raw data from all 3 DT clonal progenies were pooled and compared with pooled raw data of DS clonal progenies and commercial clones separately. Raw data were statistically analysed by two-way ANOVA by fitting a mixed effects model using GraphPad Prism 9 software (GraphPad, CA, US). Tukey’s multiple comparisons test was performed with individual variances for each comparison. Differences were considered to be statistically significant when *P* ≤ 0.05.

## Results

To investigate whether field-grown DT clones have the ability to successfully transmit drought acclimation strategies to their subsequent clonal progenies, we attempted to propagate five vegetative cuttings from three DT and DS clones at Yalumba Nursery (Nuriootpa, SA, Australia). Even though, all three DT clones were successful propagated, only two DS clones were regenerated successfully.

### Variations in soil moisture depletion during two dehydration cycles

In order to identify drought stress responses of dry-farmed clonal progenies, vines in the treatment group were subjected to two recurrent drought cycles in the glasshouse (Figure 1a). Once irrigation was withheld, the soil moisture content progressively decreased in all potted vines during D1 cycle. In comparison, a steeper decline was observed during the second drought cycle leading up to D2 (Figure 1a, b, c). This rapid reduction of soil moisture leading to D2 could be attributed to increased vapour pressure deficit (VPD) caused by slightly higher day/night temperature and marked reduction in relative humidity (Figure 1d, Table 1). Soil moisture content decreased more rapidly in commercial clones relative to other clones at D1, and DS clones maintained the highest soil moisture (Figure 1b). However, no marked differences were observed between clones during the second drought event (Figure 1c).

**Table 1:**
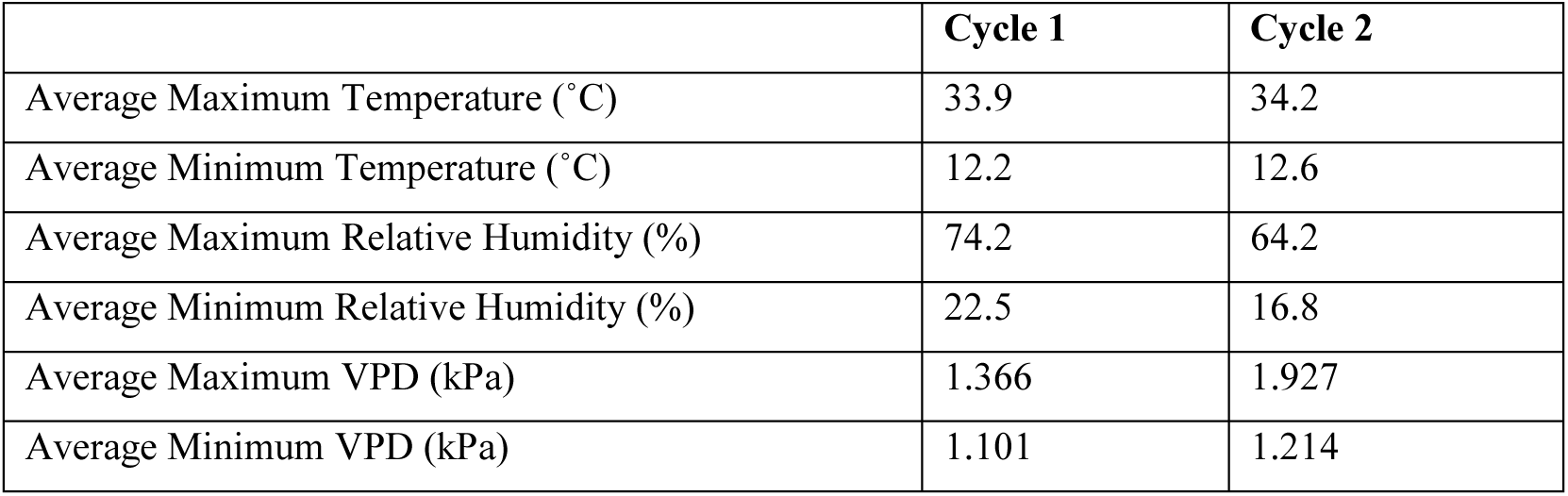
Glasshouse environmental conditions during two drought cycles.

|  | Cycle 1 | Cycle 2 |
| --- | --- | --- |
| Average Maximum Temperature (°C) | 33.9 | 34.2 |
| Average Minimum Temperature (°C) | 12.2 | 12.6 |
| Average Maximum Relative Humidity (%) | 74.2 | 64.2 |
| Average Minimum Relative Humidity (%) | 22.5 | 16.8 |
| Average Maximum VPD (kPa) | 1.366 | 1.927 |
| Average Minimum VPD (kPa) | 1.101 | 1.214 |

### Effect of differential mid-day stem water potential (Ψ_s_) and gas exchange on photosynthetic performances of dry-farmed clonal progenies under multiple drought events

In the control group, all irrigated clones had *Ψs* ∼ −0.4 MPa prior to drought stress for cycle 1, but prior to cycle 2, *Ψ*_*s*_ was significantly lower, between −0.6 & −0.8 MPa (Figure 2a). In drought-treated vines, *Ψ*_*s*_ was drastically reduced to −1.2 and −1.1 MPa at D1 and D2, respectively, indicating a moderate to high drought stress condition, with similar *Ψ*_*s*_ values between clones. None of the clones showed wilting symptoms or chlorosis of leaves even under these low *Ψ*_*s*_ values. All clones recovered after rewatering (Figure 2a).

**Figure 2.**
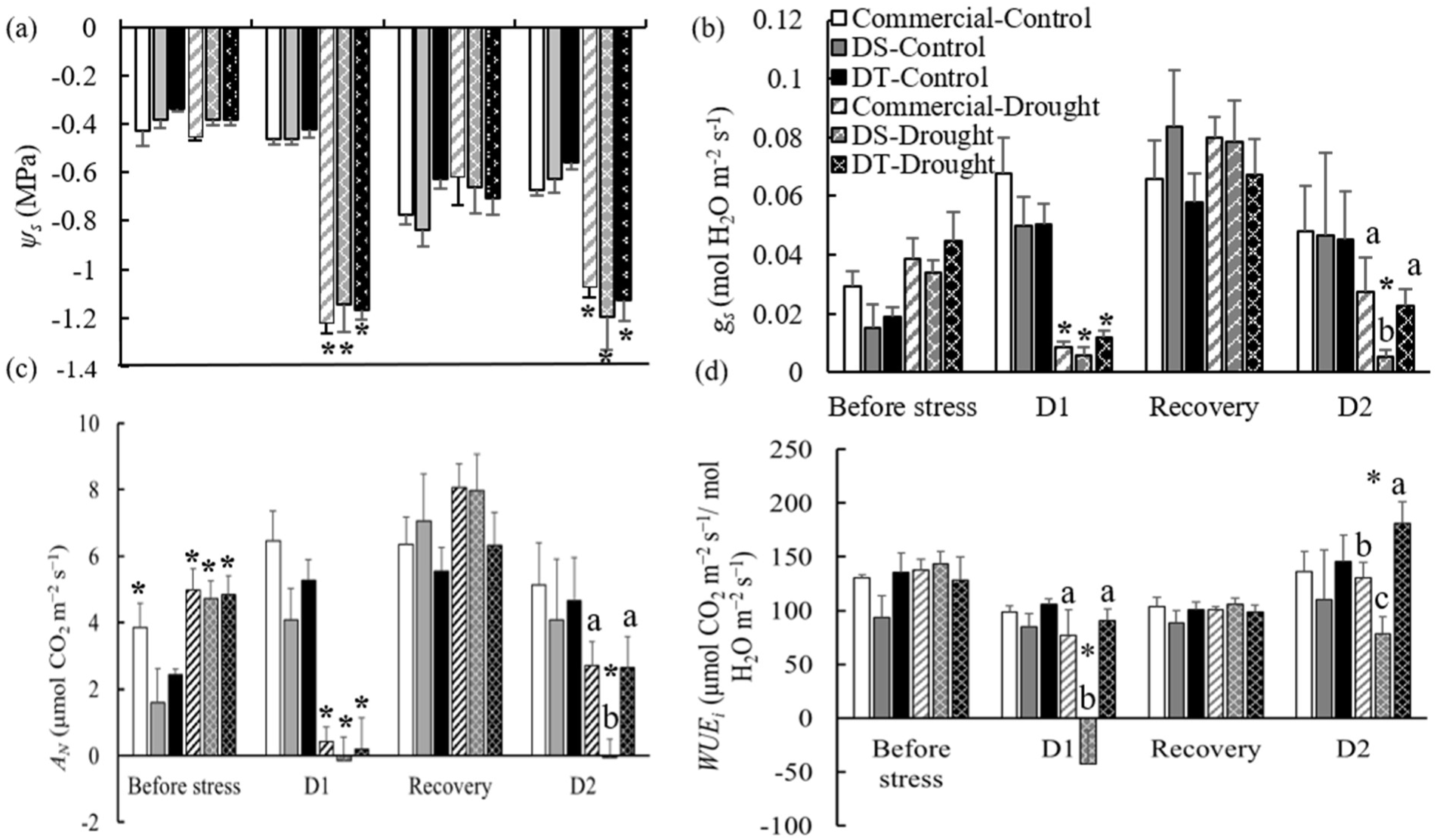
Variations in mid-day stem water potential (*Ψ*_*s*_; a), stomatal conductance (*g*_*s*_; b), net carbon assimilation (*A*_*N*_; c), and instantaneous water use efficiency (*WUE*_*i*_; d) in commercial, DS and DT clones before imposition of the stress, at the first drought cycle (D1), after rewatering and at the second drought cycle (D2). Values are means ± SEM of 4-9 biological replicates. Statistical analysis was conducted using two-way ANOVA. Asterisks indicate statistically significant differences (*P*<0.05) between drought-treated clones and controls at each treatment. Lowercase letters indicate statistically significant differences (*P*<0.05) between drought treated clones within each treatment.

In order to understand stomatal responses to soil drying, leaf gas exchange parameters were analysed in all clones under drought conditions. It is important to note that, well-watered vines as well as drought treated vines before commencement of the drought stress, displayed low *g*_*s*_ values which ranged from 0.02 to 0.08 mol H_2_O m^-2^ s^-1^ and low *A*_*N*_ (from 2 to 7 μmol CO_2_ m^-2^ s^-1^) irrespective of abundant soil moisture (Figure 2b, c). As those vines vigorously grew without any sign of water stress (leaf chlorosis or root growth restrictions) throughout the experiment, we speculated that low gas exchange values might be because of the low light intensities provided during vine growth and measurements (<50 and 1000 μmol m^-2^ s^-1^ respectively). Differences in *g*_*s*_ was similar between clones at D1 (Figure 2b). For instance, when clones were exposed to the first drought event, *g*_*s*_ decreased in commercial, DS and DT clones by 78%, 83% and 74%, respectively compared to the clones in the control group. All clones completely recovered upon rewatering. During the second drought cycle, DS clones showed further decline in *g*_*s*_ (93%) relative to recovery, whereas it was significantly higher in both commercial and DT clones relative to DS clones and at D1 (Figure 2b).

Even though in the control group, all well-irrigated clones exhibited similar plant water status and *g*_*s*_, it was consistently observed that *A*_*N*_ of commercial clones was significantly higher relative to other clones at early developmental stages. However, over time DS and DT clones also increased *A*_*N*_ to a similar level as commercial clones, therefore no statistical differences were apparent between clones at the latter developmental stages (Figure 2c). At D1, *A*_*N*_ steeply decreased to near zero in all drought-stressed clones. Although *A*_*N*_ of DS clones was also drastically reduced to near zero at D2, in commercial and DT clones, reduction of *A*_*N*_ was approximately 66% and 58% respectively (Figure 2c). Leaf *WUE*_*i*_ which represents the ratio of *A*_*N*_ versus *g*_*s*_, was similar in all clones before imposing the drought stress (Figure 2d). At the first drought cycle, *WUE*_*i*_ was marginally reduced in commercial and DT clones by 44% and 29% respectively, but 83% significant reduction was observed in DS clones. All clones displayed basal level of *WUE*_*i*_ upon rewatering. When the recovered vines were exposed to the second drought event, commercial clones exhibited a 29% increase in *WUEi*, whereas it was marginally decreased in DS clones (26%), while, an 83% significant increase in *WUEi* was observed in DT clones. Intriguingly, DT clones exhibited significantly higher *WUE*_*i*_ relative to both commercial and DS clones at D2 (Figure 2d).

### Variations in mesophyll conductance and chlorophyll fluorescence parameters in dry-farmed Cabernet clonal progenies under prolonged drought stress

In comparison to the previously reported *g*_*m*_ values (0.1-0.15 mol CO_2_ m^-2^ s^-1^) in Cabernet Sauvignon [21], 68% reduction of *g*_*m*_ was observed in all clones at D1, but when the second drought stress was imposed, it was further depleted in both commercial and DS clones by 82% and 84% respectively (Figure 3a). In contrast, *g*_*m*_ values in DT clones were significantly higher relative to other two clones at D2 (Figure 3a). Drought stress provokes remarkable changes in the photosystem II activity and chlorophyll fluorescence traits. Under non-stressed conditions, the maximum efficiency of the photosystem II (PSII) photochemistry (*F*_v_*/F*_m_) and fraction of open PSII reaction centres (*qP*) have reported to be around 0.75-0.80 [40,50] in Cabernet Sauvignon. Additionally, thermal dissipation of excess light energy (non-photochemical quenching; *NPQ*) and actual photochemical efficiency of PSII (Φ_PSII_) are approximately 0.5 and 0.2, respectively [40].

**Figure 3.**
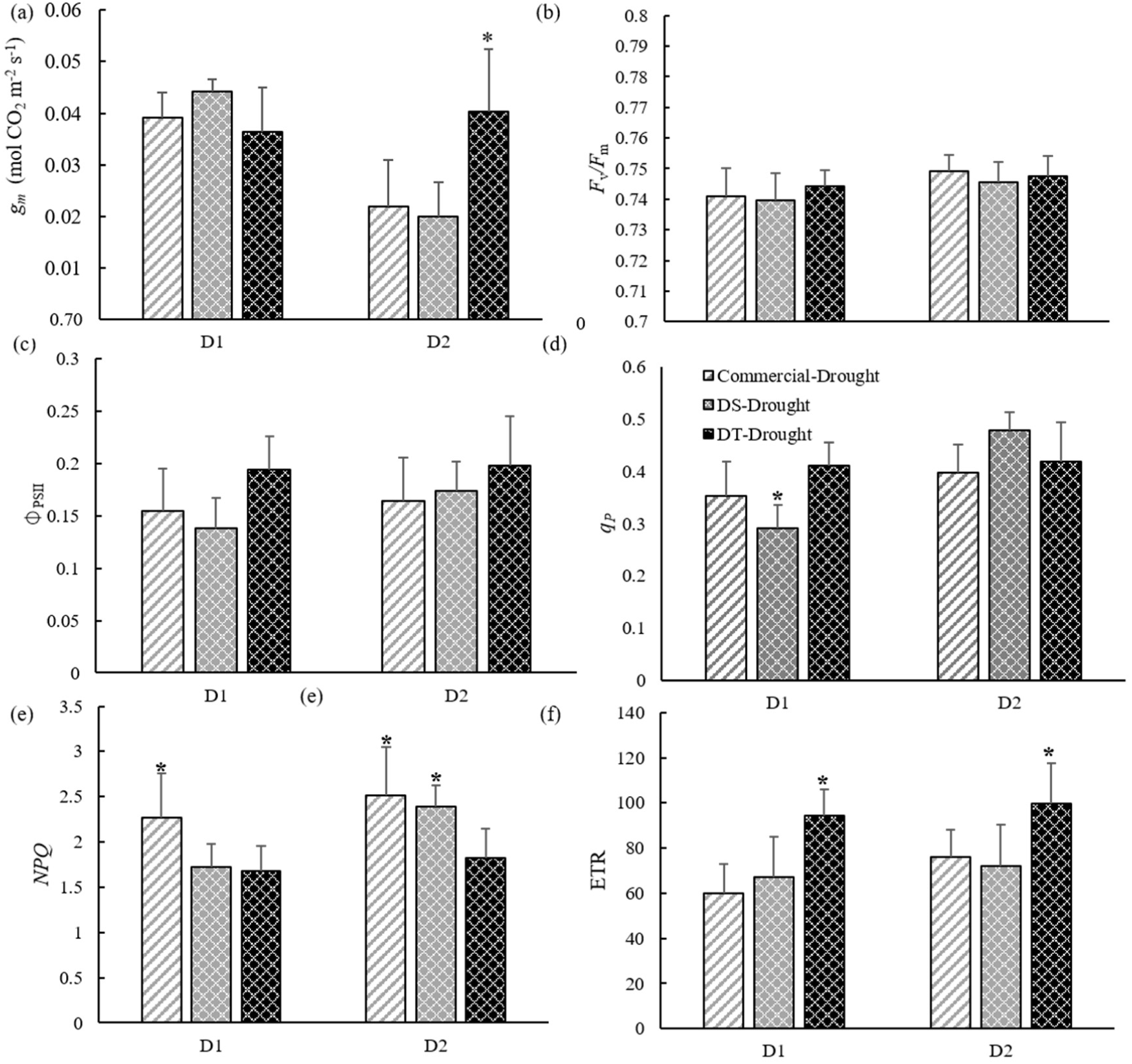
Changes in chlorophyll fluorescence parameters in dry-farmed and commercial clones at the peak of 1^st^ and 2^nd^ drought cycles. (a) Mesophyll conductance (*g*_*m*_), (b) maximum photochemical efficiency (F_v_/F_m_), (c) actual photochemical efficiency (Φ_PSII_), (d) photochemical quenching coafficient (*qP*), (e) non-photochemical quenching (*NPQ*), and (f) electron transport rate (ETR). Values are means ± SEM of six biological replicates. Statistical analysis was conducted using two-way ANOVA. Asterisks indicate statistically significant differences (*P*<0.05) between clones within each drought cycle.

In comparison to those reported values, in our study, *F*_v_*/F*_m_ remained unchanged at 0.74-0.75 in all clones at both D1 and D2, and differences could not be detected between clones (Figure 3b). Marginal decrease in Φ_PSII_ was observed only in DS and commercial clones upon exposure to both drought events, but in DT clones Φ_PSII_ remained unchanged at 0.2 (Figure 3c). All clones demonstrated substantial decline in *qP* at both drought cycles. For instance, at D1, DS clones exhibited a marked reduction in *qP* (62.6%) (Fig. 3d), whereas both commercial and DT clones displayed only 54.7% and 48.8% decline respectively. *qP* was similar between clones at D2 (Figure 3d). Dissipation of excess thermal energy within chlorophyll-containing complexes via non-photochemical quenching (*NPQ*), helps prevent the likelihood of formation of ROS [51]. During both drought cycles, *NPQ* was significantly higher in all clones relative to non-stressed conditions [40] (Figure 3e). During the first drought cycle, commercial clones displayed significantly higher *NPQ*, relative to dry-farmed clones. At the second drought stress, increase in *NPQ* was also observed in DS clones, however in DT clones *NPQ* remained at relatively lower level. It was interesting to note that electron transport rate (ETR) was significantly higher in DT clones relative to commercial and DS clones at both drought events (Figure 3f).

### Water stress-induced changes in expression of AQPs (VvTIP2;1 and VvPIP1;1) in leaves

To understand inter-clonal variation in aquaporin-mediated hydraulic regulatory mechanisms, we analysed expressions of AQP genes encoding a tonoplast intrinsic protein, VvTIP2;1, and a plasma membrane intrinsic protein, VvPIP1;1 under drought stress conditions. Aquaporin transcript abundance differed significantly between clones in response to dehydration. For instance, before clones were exposed to drought stress, *VvTIP2;1* transcript abundance was significantly higher in both DS and DT clones relative to commercial clones. Irrespective of the treatment, commercial clones exhibited basal level of *VvTIP2;1* expression. However, at D1, *VvTIP2;1* expression was significantly down-regulated in both DS and DT clones and it remained constant in both clones at recovery. At D2, the transcript abundance of *VvTIP2;1* in DS clones was significantly lower relative to both commercial and DT clones (Figure 4a). *VvPIP1;1* expression was similar in all clones before exposure to drought stress whereas it was slightly down-regulated in both commercial and DT clones at D1. However, significant up-regulation of *VvPIP1;1* was observed in DS clones at D1. At the recovery stage, both commercial and DS clones displayed basal level of *VvPIP1;1* expression, but DT clones exhibited statistically non-significant increase in *VvPIP1;1* expression. At D2, *VvPIP1;1* transcript abundance was markedly increased in commercial clones. In contrast, it was decreased in both DS and DT clones at D2 (Figure 4b).

**Figure 4.**
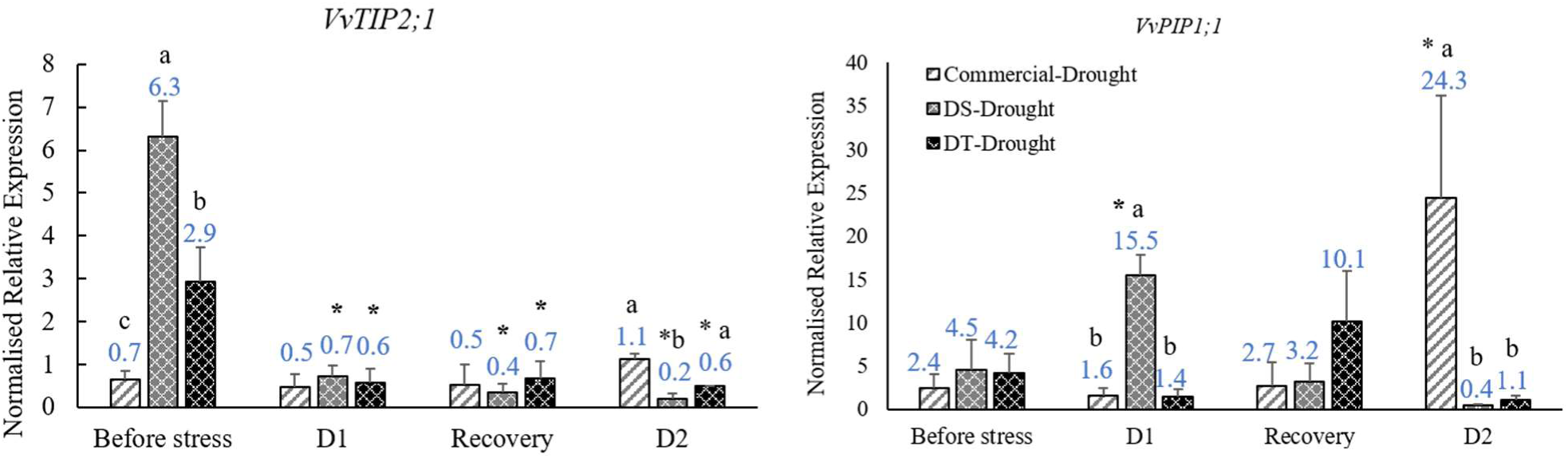
Relative gene expression (fold changes) of two aquaporin genes, a) *VvTIP2;1* and b) *VvPIP1;1* in Cabernet clones in response to dehydration and rehydration during two drought cycles. Values are means ± SEM of four biological replicates. Statistical analysis was conducted using two-way ANOVA. Asterisks indicate statistically significant differences (*P*<0.05) of drought treated clones relative to before stress. Lowercase letters indicate statistically significant differences (*P*<0.05) between drought treated clones within each treatment.

### Drought-mediated changes in ROS detoxification

To investigate whether exposure to prolonged drought stress enhances antioxidative mechanisms in Cabernet clones, ascorbate peroxidase (APX) enzyme activity were evaluated. It is interesting to note that APX activity increased significantly in commercial and DS clones at D1, but it did not change in DT clones at both drought events. Upon rewatering, APX activity remained higher in commercial clones, however both DS and DT clones displayed basal level of APX activity. At the second drought event, commercial and DT clones exhibited basal level of APX enzyme activity at D2. Unfortunately, APX activity in DS clones could not be reliably detected at D2 due to sample cross-contamination (Figure 5).

**Figure 5.**
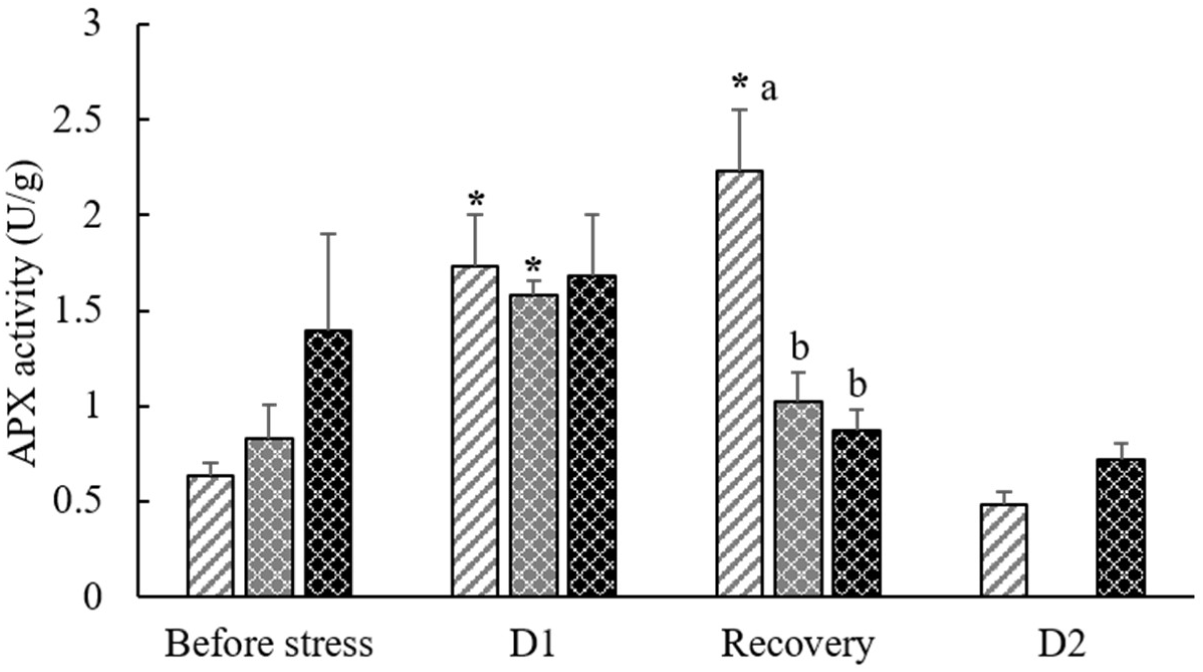
Changes in APX enzyme activity in Cabernet clones under two recurrent drought cycles. Values are means ± SEM of six biological replicates. Statistical analysis was conducted using two-way ANOVA. Asterisks indicate statistically significant differences (*P*<0.05) of drought treated clones relative to before stress. Lowercase letters indicate statistically significant differences (*P*<0.05) between drought treated clones within each treatment.

### Variations in ABA in the xylem sap (ABA_xyl_) and GABA accumulation in leaf tissues

As large body of evidence has demonstrated that significant increase in ABA in leaves is highly correlated with the abundance of ABA_xyl_, we investigated the drought-mediated changes in ABA_xyl_ in all Cabernet clones. ABA_xyl_ concentration increased significantly in all clones during both drought treatments and no marked differences were observed between clones. It returned to the basal levels upon rewatering (Figure 6a). Before imposition of the drought stress, similar GABA levels were detected in all three clones. In line with previous studies [28,29,34,35], 2-fold and 12-fold increase in GABA concentration was observed in DS and DT clones respectively at D1. However, in commercial clones, leaf GABA levels did not change in response to soil moisture content. After rewatering and at D2, basal GABA concentrations were detected in all clones (Figure 6b).

**Figure 6.**
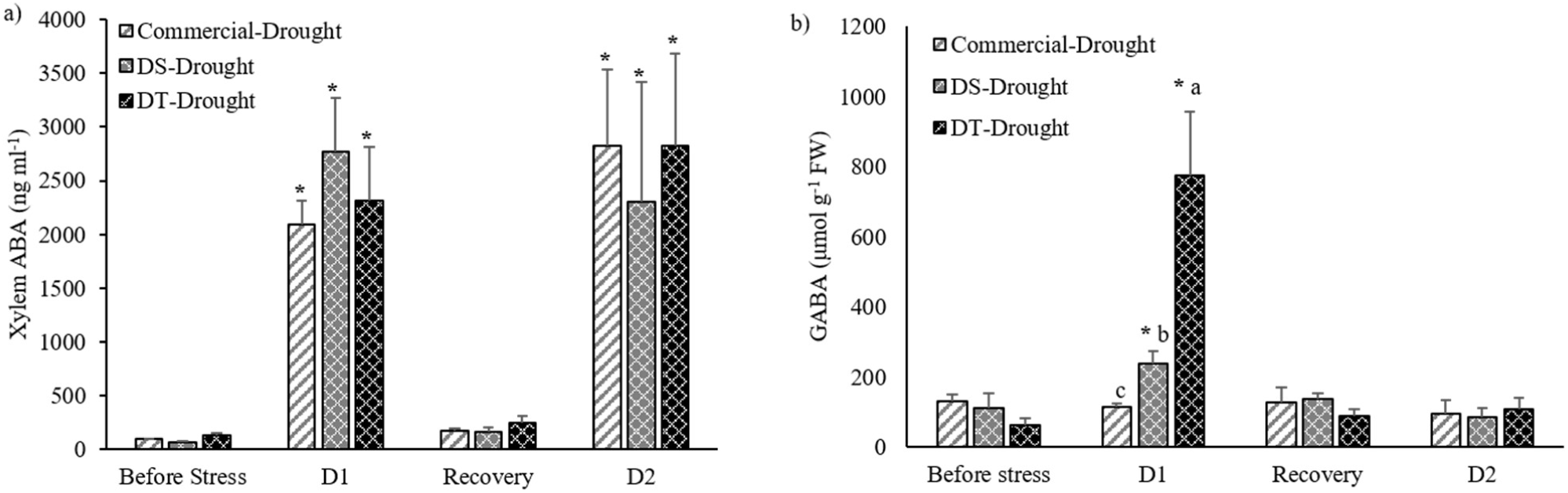
Variations in a) ABA concentrations in the xylem sap (ABA_xyl_) and, b) GABA concentrations in leaves of Cabernet clones upon exposure to two dehydration events. Values are means ± SEM of six biological replicates. Statistical analysis was conducted using two-way ANOVA. Asterisks indicate statistically significant differences (*P*<0.05) of clones relative to before exposure to drought stress. Lowercase letters indicate statistically significant differences (*P*<0.05) between drought treated clones within each treatment.

## Discussion

As for many commercial grapevine clones, unique range of viticultural and oenological traits could exist between Cabernet dry-farmed clones used in our study due to an accumulation of somatic mutations over long period of time. However, so far, no studies have been conducted to explore the genetic variations that underlies their phenotypic differences. Previously we reported that all superior shallow-rooted DT clones, identified through our preliminary field trial exhibited significantly higher *WUE*_*i*_ under limited soil moisture relative to all selected deep-rooted DS clones grown with more soil moisture [38]. This observation tempted us to speculate that irrespective of their potential individual genotypic and phenotypic diversity, all DT clones may have primed in a still unknown mechanism to perform better under limited soil moisture than all DS clones. In order to test this hypothesis, all DT, DS and commercial clonal progenies were grouped separately and their physiological and molecular responses were evaluated under two recurrent cyclical drought conditions. In support of our hypothesis, Zamorano *et al*. (2021) [24] recently demonstrated that previous season drought stress can significantly improve photosynthesis rate and *WUE*_*i*_ in field-grown grapevines under recurrent drought events. Findings of our glasshouse study further confirm the fact that field-grown DT clones have a greater ability to transmit *WUE*_*i*_ and other drought adaptation mechanisms to their subsequent clonal progenies under recurrent cyclical drought episodes relative to DS clones. In our study, transgenerational drought adaptation capability of the progeny of field-grown grapevine clones is reflected at the first drought cycle whereas their intragenerational adaptation is represented under the second drought event. Based on our findings, we propose simplified models which represents differential molecular and physiological mechanisms underpinning transgenerational and intragenerational drought stress priming in DT and DS clonal progenies and commercial clones (Figure 6).

### Differential water transport capacities of grapevine clones under drought stress

During the first drought cycle, even though all clones displayed similar water flux through stomata (*g*_*s*_) and *Ψ*_*s*_, soil moisture content depleted more rapidly in commercial clones relative to DT and DS clones (Figure 1b, 2a, 2b). At the second drought event, all clones were able to maintain similar *Ψ*_*s*_ irrespective of the higher evaporative demand (Figure 1c, 2a, 2b). Given that stomatal regulation of transpiration has a positive correlation with leaf hydraulic conductance (*K*_leaf_) [52] and root hydraulic conductivity (*Lp*_*r*_) [53,54], this observation tempted us to speculate that commercial clones may have higher *K*_leaf_ and root water uptake capacity (*K*_root_) to facilitate higher water transport than DT and DS clones.

Whole-plant water transport occurs through three routes i.e., apoplastic (through cell walls), symplastic (through plasmadesmata) and transcellular (across cell membrane) [54]. Some of the AQP isoforms belonging to the major intrinsic family (MIP) are considered as the key intra/intercellular water and ion channels which regulate transcellular or radial water transport in both leaves and roots [47,55,56]. The function and regulation of AQP are highly variable among distinct isoforms [47,54]. Previous studies have shown that water permeability of VvTIPs is higher than VvPIPs particularly to allow the cells to recruit the vacuolar space for water storage [53,56]. As *VvTIP2;1* expression in leaves was found to be well correlated with *g*_*s*_ and *K*_leaf_ [57], we examined its expression patterns in grapevine leaf tissues. Additionally, changes in transcript abundance of *VvPIP1;1* was also analysed because *VvPIP1;1* expression positively correlates with *K*_leaf_ in isohydric grapevine varieties.

In line with previous studies [47,57], *VvTIP2;1* expression pattern seems to have a close association with *g*_*s*_ under two drought events. Significant upregulation of *VvPIP1;1* was detected in DS and commercial clones at D1 and D2 respectively and DT clones exhibited statistically non-significant increase at the recovery (Figure 4a). In line with our findings, previous studies have also shown that *VvPIP1;1* expression significantly increases [47,58] or stably expressed [57] in different grapevine varieties under drought stress. Collectively, our study suggests that at D1, AQP-driven K_leaf_ is likely to be downregulated in DT clones, and upregulated in DS clones while commercial clones maintain constant *K*_leaf_. At the second drought stress, *K*_leaf_ may have been upregulated in commercial clones and down-regulated in both DT and DS clonal progenies. Higher soil moisture content in DT clones relative to commercial clones under two drought events can be explained by the reduction of water uptake. As isohydric varieties such as Cabernet Sauvignon are highly vulnerable to embolism, reduction of *K*_leaf_ has been proposed as a favourable mechanism to prevent building up the xylem tension which leads to cavitation [59,60]. Significantly higher and lower soil moisture percentages in DS and commercial clones relative to other clones at D2 can primarily be attributed to changes in plant hydraulics. As AQP expression does not directly correlate with the soil drying patterns of DS and commercial clones at D1, we speculate that differences in leaf area might have influenced their soil drying rates under drought stress. However, future studies are needed to confirm the exact roles of AQP in regulating *Lp*_*r*_ in dry-farmed grapevine clones. Collectively, our findings suggest that DT clones may have adapted to consume less amount of water upon dehydration, whereas commercial clones may require relatively higher amount of water for maintaining plant metabolism under water stress.

### Differential stomatal-regulatory mechanisms exist in dry-farmed and commercial grapevine clones under drought

During our experimental conditions, all clones exhibited near zero *g*_*s*_at D1. In response to second drought event, *g*_*s*_was further declined in DS clones whereas DT and commercial clones showed significant improvement in *g*_*s*_ (Figure 2b). In order to understand whether differential stomatal responses among clones is related to different sensitivities to key chemical and hydraulic stomatal mediators, we examined changes in concentrations of ABA_xyl_and GABA in leaf tissues in addition to *AQP* expression under two drought events.

In line with previous studies [61,62], ABA_xyl_was significantly increased in all clones during both drought events (Figure 6a). It has long been known that ABA induces stomatal closure either directly via the activation of ion channels or indirectly via restricting radial water flow from the xylem by down-regulating AQP expression [63,64]. Given the changes in AQP-mediated *K*_leaf_and GABA concentrations in leaves, it can be proposed that at D1, stomatal closure in DS clones might have induced by an additive effects of chemical (ABA- and GABA-driven) mechanisms whereas both chemical and hydraulic mechanisms are likely to be exist in DT clones (Figure 7). At D2, even though both ABA- and AQP-mediated stomatal regulatory mechanisms seem to exist in both DS and DT clones, significant reduction in *g*_*s*_ was observed only in DS clones. Interestingly, commercial clones do not seem to have hydraulic and GABA-dependant stomatal regulation under both drought events. Even though, increased accumulation of ABA_xyl_has shown to downregulate *g*_*s*_ in commercial clones at D1, similar *g*_*s*_ reduction was not detected at D2 (Figure 2b). Therefore, further investigations are required to understand whether there is another internal stimuli which prevents stomatal closure in both DT and commercial clones under recurrent drought events (Figure 7). For instance, recent work in *Arabidopsis* have shed light on CLAVATA3/embryo-surrounding region-related (CLE) peptides as novel messengers which are involved in root-to-shoot signalling for regulating stomatal aperture movements under drought [18,65]. However, no detailed investigations have been conducted in grapevine to understand their specific functions in stomatal regulation under long-term drought stress.

**Figure 7.**
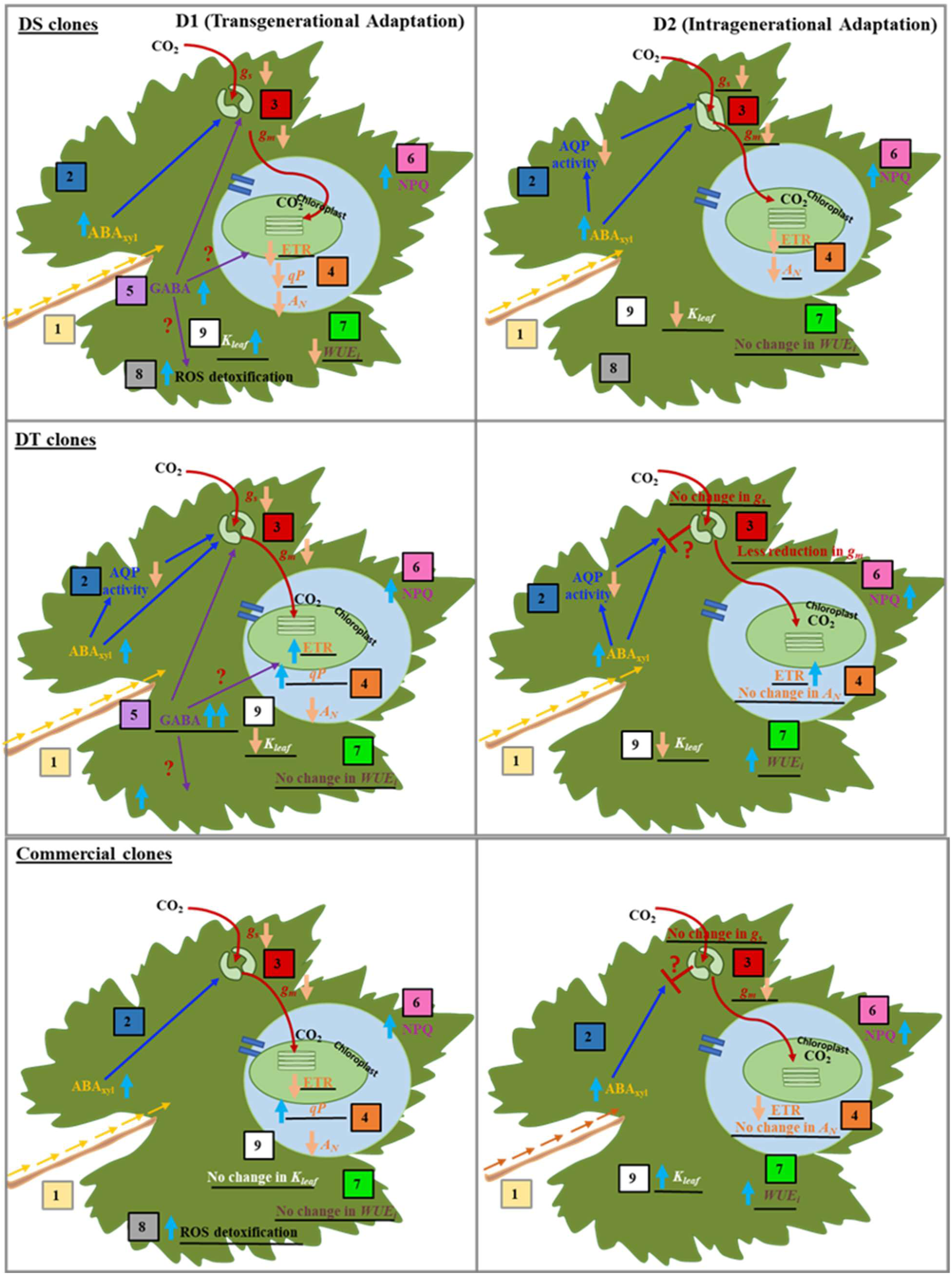
Schematic representation of differential long-term drought acclimation mechanisms of DS (top), DS (middle) and commercial (bottom) clones during first (D1; left) and second (D2; right) drought cycles. The diagram represents 1) accumulation of ABA_xyl_ in the leaf (yellow arrows and letter), 2) ABA-dependent or ABA-AQP-mediated stomatal regulatory mechanisms (blue arrows and letter), 3) CO_2_ influx through stomatal and mesophyll conductance (red arrows and letters), 4) photosynthesis-associated processes (orange letters), 5) GABA-mediated priming responses (purple arrows and letters), 6) non-photochemical quenching (pink letters), 7) *WUE*_*i*_ (brown letters), 8) antioxidative defence responses (black letters) and, 9) leaf hydraulic conductance (*K*_leaf_; white letters). Differential physiological and molecular responses observed between clones at each drought event have been underlined. Light blue thick arrow indicates an increase or upregulation whereas orange arrow represents a decrease or downregulation. Our findings support a model in which DT clones exhibits a more efficient transgenerational drought acclimation mechanisms relative to DS clones through GABA-mediated priming and improving photochemical efficiency and *WUE*_*i*_ at D1. Additionally, improvement in CO_2_ influx through stomatal conductance, photochemical efficiency and *WUE*_*i*_ also contributes to intragenerational stress priming of DT and commercial clones relative to DS clones at D2. Red question marks denote mechanistic points warranting further investigations.

### Effect of differential hydraulic, stomatal and non-stomatal regulatory mechanisms on photosynthetic performance of dry-farmed clones

Our findings suggest that DT and commercial clones are able to improve plant photosynthetic performances to adapt faster to prolonged drought episodes than DS clones (Figure 2b). Overall, our results indicate that the significant decline in *A*_*N*_ in all clones at D1 and in DS clones and D2 are likely to be associated with a decrease in *g*_*s*_ and *qP*. Interestingly, under prolong drought stress, improved photosynthetic performances of both commercial and DT clones can be explained as a result of improved CO_2_ diffusion and activation of their photoprotective mechanisms.

Under steady state conditions, major metabolic processes such as photosynthesis generate highly toxic reactive oxygen species (ROS), but the potential cytotoxicity is minimized by activating ROS detoxification mechanisms. However, when plants are exposed to drought, the delicate equilibrium between ROS production and scavenging is perturbed due to limitations on CO_2_ assimilation [66]. Different grapevine varieties possess various photoprotective mechanisms to cope with drought-mediated photoinhibition [67,68]. For instance, enhanced electron transport rate (ETR) facilitates channelling of majority of excess electrons to photosystems [40,42]. Additionally, the absorbed excess light energy can be dissipated as heat through non-photochemical quenching (*NPQ*) [69]. Comprehensive studies demonstrating intervarietal clonal differences in photo-protective mechanisms are still scarce. Our study demonstrate that such diversity is still exist between grapevine clones upon dehydration.

Previous studies have shown that APX plays a vital role in removing hydrogen peroxide (H_2_O_2_), which is the primary photosynthesis-associated ROS [70]. In line with previous studies, APX activity was significantly increased only in both commercial and DS clones at D1, but DT clones did not show increase in APX activity under both drought cycles (Figure 5). Significantly lower *NPQ* and higher ETR in DT clones at both drought events also imply that DT clones may have low level of acute oxidative stress relative to other clones (Figure 3e, f). As previous studies have shown that in mitochondria GABA is catabolised into Succinate, which acts as an electron donor to the mitochondrial electron transport, we speculated that increased GABA accumulation in DT clones may help redirecting excess photochemical energy through mitochondrial electron transport system to enhance cellular respiration and therefore, DT clones may have reduced ROS-mediated photo-oxidative damages at D1 relative to both DS and commercial clones. It is interesting to further investigate whether increased electron transport in the light harvesting complex is also mediated by GABA. Even though, commercial clones do not possess GABA-mediated ROS-detoxification mechanisms, they seemed to maintain cellular homeostasis by activating antioxidation system and non-photochemical quenching under drought. However, increased APX activity and *NPQ* and low ETR in DS clones would indicate that 2-fold accumulation of GABA may not be sufficient to provide complete protection against drought-induced photooxidative damages in DS clones (Figure 3e, f, 5).

The GABA is also considered as an important priming agent which allows plants to adapt faster and stronger to subsequent stress events, however, its priming-associated mechanisms are largely unknown. Given the fact that DT clones had similar level of APX activity, *NPQ* and ETR at both D1 and D2, our study in part suggests that GABA may also play a crucial role in intragenerational priming in DT clones to enhance their long-term drought resilience. However, some fundamental questions remain unanswered. If GABA is deemed bona fide priming elicitor, does it mediate stomatal regulation, photochemical efficiency and oxidative stress tolerance in grapevine clones? (Fig. 6) Are epigenetic modifications such as DNA methylation, histone modification or chromosomal remodelling also contribute to stress priming? Our future studies will elucidate whether long-term drought adaptation of DT clones is caused by genetic or epigenetic modifications. Ultimately, the fundamental insights obtained from this study will pave the way for future research aiming towards understanding whether we can “stress-memory”-associated candidate genes/epigenes to develop primed-grapevine varieties for a range of changing environments. Additionally, this study will also be beneficial for the grapevine industry, to develop/select drought resilient genotypes and clones suitable for grapevine breeding programs.

## Acknowledgements

The authors would like to thank Dr. Catherine M. Kidman at Wynns Coonawarra Estate and Yalumba Nursery for providing plant materials and their in-kind support and Annette Betts for assisting with the ABA analyses.

## References

1. Report, W.A. Economic Contribution of the Australian Wine Sector 2019; Wine Australia: 2019.

2. Report, N.V. National Vintage report 2020_WA. 2020.

3. IPCC. Climate change 2021 the physical science basis; 7/08/2021 2021.

4. Nicholas, P. Grapevine clones used in Australia; South Australian Research and Development Institute: 2006.

5. Vondras, A.M.; Minio, A.; Blanco-Ulate, B.; Figueroa-Balderas, R.; Penn, M.A.; Zhou, Y.; Seymour, D.; Ye, Z.; Liang, D.; Espinoza, L.K. The genomic diversification of grapevine clones. BMC genomics 2019, 20, 1–19.

6. Xie, H.; Konate, M.; Sai, N.; Tesfamicael, K.G.; Cavagnaro, T.; Gilliham, M.; Breen, J.; Metcalfe, A.; Stephen, J.R.; De Bei, R. Global DNA methylation patterns can play a role in defining terroir in grapevine (Vitis vinifera cv. Shiraz). Frontiers in plant science 2017, 8, 1860.

7. Varela, A.; Ibañez, V.N.; Alonso, R.; Zavallo, D.; Asurmendi, S.; Talquenca, S.G.; Marfil, C.F.; Berli, F.J. Vineyard environments influence Malbec grapevine phenotypic traits and DNA methylation patterns in a clone-dependent way. Plant Cell Reports 2021, 40, 111–125.

8. Atak, A.; Kahraman, K.; Söylemezoğlu, G. Ampelographic identification and comparison of some table grape (Vitis vinifera L.) clones. New Zealand Journal of Crop and Horticultural Science 2014, 42, 77–86.

9. Roach, M.J.; Johnson, D.L.; Bohlmann, J.; van Vuuren, H.J.; Jones, S.J.; Pretorius, I.S.; Schmidt, S.A.; Borneman, A.R. Population sequencing reveals clonal diversity and ancestral inbreeding in the grapevine cultivar Chardonnay. PLoS genetics 2018, 14, e1007807.

10. Lovisolo, C.; Lavoie-Lamoureux, A.; Tramontini, S.; Ferrandino, A. Grapevine adaptations to water stress: new perspectives about soil/plant interactions. Theoretical and Experimental Plant Physiology 2016, 28, 53–66, doi:10.1007/s40626-016-0057-7.

11. <._Lovisolo et al., 2010-drought induced Whole p¬ant hydraulic and non-hydraulic interaction in grapevine.pdf>.+

12. Coupel-Ledru, A.; Tyerman, S.D.; Masclef, D.; Lebon, E.; Christophe, A.; Edwards, E.J.; Simonneau, T. Abscisic Acid Down-Regulates Hydraulic Conductance of Grapevine Leaves in Isohydric Genotypes Only. Plant Physiol 2017, 175, 1121–1134, doi:10.1104/pp.17.00698.

13. Yoshida, R.; Umezawa, T.; Mizoguchi, T.; Takahashi, S.; Takahashi, F.; Shinozaki, K. The regulatory domain of SRK2E/OST1/SnRK2.6 interacts with ABI1 and integrates abscisic acid (ABA) and osmotic stress signals controlling stomatal closure in Arabidopsis. J Biol Chem 2006, 281, 5310–5318, doi:10.1074/jbc.M509820200.

14. Rossdeutsch, L.; Edwards, E.; Cookson, S.J.; Barrieu, F.; Gambetta, G.A.; Delrot, S.; Ollat, N. ABA-mediated responses to water deficit separate grapevine genotypes by their genetic background. BMC Plant Biol 2016, 16, 91, doi:10.1186/s12870-016-0778-4.

15. Joshi-Saha, A.; Valon, C.; Leung, J. Abscisic acid signal off the STARting block. Molecular plant 2011, 4, 562–580.

16. Kim, T.-H.; Böhmer, M.; Hu, H.; Nishimura, N.; Schroeder, J.I. Guard cell signal transduction network: advances in understanding abscisic acid, CO2, and Ca2+ signaling. Annual review of plant biology 2010, 61, 561–591.

17. Holbrook, N.M.; Shashidhar, V.; James, R.A.; Munns, R. Stomatal control in tomato with ABA-deficient roots: response of grafted plants to soil drying. Journal of Experimental Botany 2002, 53, 1503–1514.

18. Takahashi, F.; Suzuki, T.; Osakabe, Y.; Betsuyaku, S.; Kondo, Y.; Dohmae, N.; Fukuda, H.; Yamaguchi-Shinozaki, K.; Shinozaki, K. A small peptide modulates stomatal control via abscisic acid in long-distance signalling. Nature 2018, 556, 235–238.

19. Christmann, A.; Grill, E. Peptide signal alerts plants to drought. 2018.

20. Chaves, M.M. Effects of Water Deficits on Carbon Assimilation. Journal of Experimental Botany 1991, 42, 1–16.

21. Tomás, M.; Medrano, H.; Brugnoli, E.; Escalona, J.M.; Martorell, S.; Pou, A.; Ribas-Carbó, M.; Flexas, J. Variability of mesophyll conductance in grapevine cultivars under water stress conditions in relation to leaf anatomy and water use efficiency. Australian Journal of Grape and Wine Research 2014, 20, 272–280, doi:10.1111/ajgw.12069.

22. Flexas, J.; GalmÃ S, J.; GallÃ, A.; GulÃ As, J.; Pou, A.; Ribas-Carbo, M.; TomÃ S, M.; Medrano, H. Improving water use efficiency in grapevines: potential physiological targets for biotechnological improvement. Australian Journal of Grape and Wine Research 2010, 16, 106–121, doi:10.1111/j.1755-0238.2009.00057.x.

23. Alves, R.D.; Menezes-Silva, P.E.; Sousa, L.F.; Loram-Lourenço, L.; Silva, M.L.; Almeida, S.E.; Silva, F.G.; de Souza, L.P.; Fernie, A.R.; Farnese, F.S. Evidence of drought memory in Dipteryx alata indicates differential acclimation of plants to savanna conditions. Scientific reports 2020, 10, 1–16.

24. Zamorano, D.; Franck, N.; Pastenes, C.; Wallberg, B.; Garrido, M.; Silva, H. Improved physiological performance in grapevine (Vitis vinifera L.) cv. Cabernet Sauvignon facing recurrent drought stress. Australian Journal of Grape and Wine Research 2021.

25. Tombesi, S.; Frioni, T.; Poni, S.; Palliotti, A. Effect of water stress “memory” on plant behavior during subsequent drought stress. Environmental and Experimental Botany 2018, 150, 106–114.

26. Springer, N.M.; Schmitz, R.J. Exploiting induced and natural epigenetic variation for crop improvement. Nat Rev Genet 2017, 18, 563–575, doi:10.1038/nrg.2017.45.

27. Hagmann, J.; Becker, C.; Muller, J.; Stegle, O.; Meyer, R.C.; Wang, G.; Schneeberger, K.; Fitz, J.; Altmann, T.; Bergelson, J.; et al. Century-scale methylome stability in a recently diverged Arabidopsis thaliana lineage. PLoS Genet 2015, 11, e1004920, doi:10.1371/journal.pgen.1004920.

28. Ramesh, S.A.; Tyerman, S.D.; Xu, B.; Bose, J.; Kaur, S.; Conn, V.; Domingos, P.; Ullah, S.; Wege, S.; Shabala, S. GABA signalling modulates plant growth by directly regulating the activity of plant-specific anion transporters. Nature communications 2015, 6, 1–10.

29. Vijayakumari, K.; Puthur, J.T. γ-Aminobutyric acid (GABA) priming enhances the osmotic stress tolerance in Piper nigrum Linn. plants subjected to PEG-induced stress. Plant Growth Regulation 2016, 78, 57–67.

30. Bouché, N.; Lacombe, B.t.; Fromm, H. GABA signaling: a conserved and ubiquitous mechanism. Trends in Cell Biology 2003, 13, 607–610, doi:https://doi.org/10.1016/j.tcb.2003.10.001.

31. Bown, A.W.; Shelp, B.J. Plant GABA: Not Just a Metabolite. Trends in Plant Science 2016, 21, 811–813, doi:https://doi.org/10.1016/j.tplants.2016.08.001.

32. Michaeli, S.; Fromm, H. Closing the Loop on the GABA Shunt in Plants: Are GABA metabolism and signaling entwined? Frontiers in Plant Science 2015, 6.

33. Xu, B.; Long, Y.; Feng, X.; Zhu, X.; Sai, N.; Chirkova, L.; Betts, A.; Herrmann, J.; Edwards, E.J.; Okamoto, M. GABA signalling modulates stomatal opening to enhance plant water use efficiency and drought resilience. Nature communications 2021, 12, 1–13.

34. Shelp, B.J.; Bown, A.W.; McLean, M.D. Metabolism and functions of gamma-aminobutyric acid. Trends in plant science 1999, 4, 446–452.

35. Krishnan, S.; Laskowski, K.; Shukla, V.; Merewitz, E.B. Mitigation of drought stress damage by exogenous application of a non-protein amino acid γ–aminobutyric acid on perennial ryegrass. Journal of the American Society for Horticultural Science 2013, 138, 358–366.

36. Nayyar, H.; Kaur, R.; Kaur, S.; Singh, R. γ-Aminobutyric acid (GABA) imparts partial protection from heat stress injury to rice seedlings by improving leaf turgor and upregulating osmoprotectants and antioxidants. Journal of Plant Growth Regulation 2014, 33, 408–419.

37. Yang, A.; Cao, S.; Yang, Z.; Cai, Y.; Zheng, Y. γ-Aminobutyric acid treatment reduces chilling injury and activates the defence response of peach fruit. Food Chemistry 2011, 129, 1619–1622.

38. Pagay, V.; Furlan, T.S.; Kidman, C.M.; Nagahatenna, D. Long-term drought adaptation of unirrigated grapevines (Vitis vinifera L.). Theoretical and Experimental Plant Physiology 2022, doi:10.1007/s40626-022-00243-3.

39. Murchie, E.H.; Lawson, T. Chlorophyll fluorescence analysis: a guide to good practice and understanding some new applications. J Exp Bot 2013, 64, 3983–3998, doi:10.1093/jxb/ert208.

40. Guan, X.; Gu, S. Photorespiration and photoprotection of grapevine (Vitis vinifera L. cv. Cabernet Sauvignon) under water stress. 2009.

41. Rahimzadeh-Bajgiran, P.; Tubuxin, B.; Omasa, K. Estimating Chlorophyll Fluorescence Parameters Using the Joint Fraunhofer Line Depth and Laser-Induced Saturation Pulse (FLD-LISP) Method in Different Plant Species. Remote Sensing 2017, 9, doi:10.3390/rs9060599.

42. Flexas, J.; Escalona, J.; Medrano, H. Down-regulation of photosynthesis by drought under field conditions in grapevine leaves. Functional Plant Biology 1998, 25, 893–900.

43. Maxwell, K.; Johnson, G.N. Chlorophyll fluorescence—a practical guide. Journal of experimental botany 2000, 51, 659–668.

44. Galle, A.; Florez-Sarasa, I.; Thameur, A.; de Paepe, R.; Flexas, J.; Ribas-Carbo, M. Effects of drought stress and subsequent rewatering on photosynthetic and respiratory pathways in Nicotiana sylvestris wild type and the mitochondrial complex I-deficient CMSII mutant. J Exp Bot 2010, 61, 765–775, doi:10.1093/jxb/erp344.

45. Sun, J.; You, X.; Li, L.; Peng, H.; Su, W.; Li, C.; He, Q.; Liao, F. Effects of a phospholipase D inhibitor on postharvest enzymatic browning and oxidative stress of litchi fruit. Postharvest Biology and Technology 2011, 62, 288–294, doi:10.1016/j.postharvbio.2011.07.001.

46. Ramesh, S.A.; Kamran, M.; Sullivan, W.; Chirkova, L.; Okamoto, M.; Degryse, F.; McLaughlin, M.; Gilliham, M.; Tyerman, S.D. Aluminum-activated malate transporters can facilitate GABA transport. The Plant Cell 2018, 30, 1147–1164.

47. Shelden, M.C.; Vandeleur, R.; Kaiser, B.N.; Tyerman, S.D. A Comparison of Petiole Hydraulics and Aquaporin Expression in an Anisohydric and Isohydric Cultivar of Grapevine in Response to Water-Stress Induced Cavitation. Front Plant Sci 2017, 8, 1893, doi:10.3389/fpls.2017.01893.

48. Vandesompele, J.; De Preter, K.; Pattyn, F.; Poppe, B.; Van Roy, N.; De Paepe, A.; Speleman, F. Accurate normalization of real-time quantitative RT-PCR data by geometric averaging of multiple internal control genes. Genome biology 2002, 3, 1–12.

49. Burton, R.A.; Shirley, N.J.; King, B.J.; Harvey, A.J.; Fincher, G.B. The CesA gene family of barley. Quantitative analysis of transcripts reveals two groups of co-expressed genes. Plant physiology 2004, 134, 224–236.

50. Min, Z.; Li, R.; Chen, L.; Zhang, Y.; Li, Z.; Liu, M.; Ju, Y.; Fang, Y. Alleviation of drought stress in grapevine by foliar-applied strigolactones. Plant Physiol Biochem 2019, 135, 99–110, doi:10.1016/j.plaphy.2018.11.037.

51. Roach, T.; Krieger-Liszkay, A. Regulation of photosynthetic electron transport and photoinhibition. Current Protein and Peptide Science 2014, 15, 351–362.

52. Sack, L.; Holbrook, N.M. Leaf hydraulics. Annu. Rev. Plant Biol. 2006, 57, 361–381.

53. Maurel, C.; Simonneau, T.; Sutka, M. The significance of roots as hydraulic rheostats. Journal of experimental botany 2010, 61, 3191–3198.

54. Vandeleur, R.K.; Sullivan, W.; Athman, A.; Jordans, C.; Gilliham, M.; Kaiser, B.N.; Tyerman, S.D. Rapid shoot-to-root signalling regulates root hydraulic conductance via aquaporins. Plant, Cell & Environment 2014, 37, 520–538.

55. Shelden, M.C.; Howitt, S.M.; Kaiser, B.N.; Tyerman, S.D. Identification and functional characterisation of aquaporins in the grapevine, Vitis vinifera. Functional Plant Biology 2009, 36, 1065–1078.

56. Sabir, F.; Zarrouk, O.; Noronha, H.; Loureiro-Dias, M.C.; Soveral, G.; Gerós, H.; Prista, C. Grapevine aquaporins: Diversity, cellular functions, and ecophysiological perspectives. Biochimie 2021.

57. Pou, A.; Medrano, H.; Flexas, J.; Tyerman, S.D. A putative role for TIP and PIP aquaporins in dynamics of leaf hydraulic and stomatal conductances in grapevine under water stress and re-watering. Plant Cell Environ 2013, 36, 828–843, doi:10.1111/pce.12019.

58. Dayer, S.; Scharwies, J.D.; Ramesh, S.A.; Sullivan, W.; Doerflinger, F.C.; Pagay, V.; Tyerman, S.D. Comparing hydraulics between two grapevine cultivars reveals differences in stomatal regulation under water stress and exogenous ABA applications. Frontiers in plant science 2020, 11, 705.

59. Schultz, H.R. Differences in hydraulic architecture account for near-isohydric and anisohydric behaviour of two field-grown Vitis vinifera L. cultivars during drought. Plant, Cell & Environment 2003, 26, 1393–1405.

60. Gerzon, E.; Biton, I.; Yaniv, Y.; Zemach, H.; Netzer, Y.; Schwartz, A.; Fait, A.; Ben-Ari, G. Grapevine Anatomy as a Possible Determinant of Isohydric or Anisohydric Behavior. American Journal of Enology and Viticulture 2015, 66, 340–347, doi:10.5344/ajev.2015.14090.

61. Bano, A.; Dorffling, K.; Bettin, D.; Hahn, H. Abscisic acid and cytokinins as possible root-to-shoot signals in xylem sap of rice plants in drying soil. Functional Plant Biology 1993, 20, 109–115.

62. Munns, R.; Cramer, G. Is coordination of leaf and root growth mediated by abscisic acid? Opinion. Plant and Soil 1996, 185, 33–49.

63. Shatil-Cohen, A.; Attia, Z.; Moshelion, M. Bundle-sheath cell regulation of xylem-mesophyll water transport via aquaporins under drought stress: a target of xylem-borne ABA? The Plant Journal 2011, 67, 72–80.

64. Geiger, D.; Maierhofer, T.; Al-Rasheid, K.A.; Scherzer, S.; Mumm, P.; Liese, A.; Ache, P.; Wellmann, C.; Marten, I.; Grill, E. Stomatal closure by fast abscisic acid signaling is mediated by the guard cell anion channel SLAH3 and the receptor RCAR1. Science signaling 2011, 4, ra32–ra32.

65. Fletcher, J.C. Recent advances in Arabidopsis CLE peptide signaling. Trends in Plant Science 2020, 25, 1005–1016.

66. Cruz de Carvalho, M.H. Drought stress and reactive oxygen species: production, scavenging and signaling. Plant signaling & behavior 2008, 3, 156–165.

67. Hochberg, U.; Degu, A.; Fait, A.; Rachmilevitch, S. Near isohydric grapevine cultivar displays higher photosynthetic efficiency and photorespiration rates under drought stress as compared with near anisohydric grapevine cultivar. Physiol Plant 2013, 147, 443–452, doi:10.1111/j.1399-3054.2012.01671.x.

68. Guan, X.; Zhao, S.; Li, D.; Shu, H. Photoprotective function of photorespiration in several grapevine cultivars under drought stress. Photosynthetica 2004, 42, 31–36.

69. Wang, H.; Gu, M.; Cui, J.; Shi, K.; Zhou, Y.; Yu, J. Effects of light quality on CO2 assimilation, chlorophyll-fluorescence quenching, expression of Calvin cycle genes and carbohydrate accumulation in Cucumis sativus. Journal of Photochemistry and Photobiology B: Biology 2009, 96, 30–37.

70. Foyer, C.H.; Noctor, G. Redox sensing and signalling associated with reactive oxygen in chloroplasts, peroxisomes and mitochondria. Physiologia plantarum 2003, 119, 355–364.

